# Linalool and *trans*-nerolidol prevent pentylenetetrazole-induced seizures in adult zebrafish

**DOI:** 10.1101/2025.01.14.633001

**Authors:** Amanda L. Silva, Leonardo M. Bastos, Matheus Gallas-Lopes, Rafael Chitolina, Carlos G. Reis, Domingos Sávio Nunes, Elaine Elisabetsky, Ana P. Herrmann, Maria Elisa Calcagnotto, Angelo Piato

## Abstract

**Background:** Linalool (LIN) and *trans*-nerolidol (NER) are terpene alcohols found in plant-derived essential oils commonly used in traditional systems of medicine. Both compounds have shown antiseizure, sedative, and antioxidant effects in rodent and *in vitro* models. Due to their structural similarity, a comparative evaluation of their antiseizure profiles is warranted. This study examined the effects of acute LIN and NER exposure in a pentylenetetrazole (PTZ)-induced seizure model in zebrafish (*Danio rerio)*.

**Methods:** A total of 240 adult zebrafish were randomly assigned to the following groups: blank control (dechlorinated water), vehicle control (1% DMSO), positive control (50 µM diazepam, DZP), and three concentrations of LIN or NER (4, 40, 400 µM). Fish were exposed to treatments for 10 minutes, transferred to a washout beaker with dechlorinated water for 5 minutes, and then transferred to PTZ (10 mM) for 20-minute behavioral recording. Seizure activity was scored by blinded observers (BORIS®), and locomotion was analyzed with ANY-maze™ software.

**Results:** LIN at 400 µM and NER at 40 and 400 µM significantly prolonged latency to clonic- and tonic-like seizures and reduced seizure severity. Notably, 400 µM NER exceeded DZP in effect size, and 40 µM NER also enhanced locomotor activity.

**Conclusion:** LIN and NER delayed progression to severe seizure stages and reduced seizure severity, supporting their antiseizure potential. NER consistently outperformed LIN, demonstrating stronger efficacy than DZP at the highest concentration. Consistent with rodent studies, these findings position both as promising leads for drug development.

## 1. INTRODUCTION

Essential oils (EOs) are mixtures of plant-derived volatile compounds, including alcohol terpenes and phenylpropanoids (1,2). Historically, plant species that produce EOs have been used in traditional medical systems to treat neuropsychiatric conditions such as anxiety, depression, and epilepsy (2–4). For example, EOs from sour orange (*Citrus aurantium* L.) have been used as alternative therapies for anxiety, insomnia, and seizure-like symptoms in Brazilian, Chinese, and Haitian folk medicine (5,6). Another example is a well-known Amazonian homemade antiseizure remedy that includes leaves from *Aeollanthus suaveolens* (Mart. ex Spreng) (7). Studies have identified linalool as the primary active component in this remedy, with strong antiseizure effects demonstrated across a variety of rodent models (1). Interestingly, its mechanism of action—investigated both in vivo and in vitro—appears to involve the inhibition of potassium-stimulated (but not basal) glutamate release and antagonism of NMDA receptors, without significant direct interaction with the GABAergic system.

Unfortunately, up to 30% of patients with epilepsy are resistant to currently available antiseizure medications (8), underscoring the urgent need for new drugs with novel mechanisms of action. Most antiseizure medications approved by the U.S. Food and Drug Administration (FDA) work by modulating GABAergic receptors (e.g., diazepam, clonazepam, lorazepam) or voltage-gated sodium channels (e.g., carbamazepine, lamotrigine, topiramate). While the mechanism of action of many EOs is thought to involve GABAergic modulation and voltage-gated channel activity (2), the unique mode of action of linalool offers promising potential for pharmacodynamic innovation in antiseizure therapy.

Linalool (LIN) and *trans-*nerolidol (NER) are acyclic terpenoids synthesized as secondary metabolites during protein metabolism in various plant species, including rosemary (*Salvia rosmarinus L.*), lavender (*Lavandula angustifolia* Mill.), and sour orange (*Citrus aurantium* L.) (9–13). Structurally, both compounds share a linear carbon backbone and hydroxyl functional group, but differ in chain length and degree of unsaturation, which may influence their pharmacological activity. Comparing structurally similar compounds like LIN and NER can help identify key structural features responsible for specific bioactivities—an approach particularly useful in the development of novel antiseizure medications. In contrast, NER antiseizure potential has only been reported in a single kindling model study (1), limiting direct comparisons between the two compounds and leaving its broader profile largely unexplored.

Zebrafish (*Danio rerio*, Hamilton) have become an increasingly popular model organism for antiseizure drug screening. A recent systematic review identified pentylenetetrazole (PTZ) as the most commonly used chemical agent to induce seizures in zebrafish, followed by kainic acid and pilocarpine (14). As early as the 2000s, it was demonstrated that PTZ-treated zebrafish larvae exhibit chemically induced seizure-like electrical discharges and behavioral phenotypes relevant to epilepsy research (15). This model was adapted for adult zebrafish, allowing for more detailed behavioral characterization and a better understanding of seizure-related alterations caused by GABA_A_ receptor antagonism in this species (16). In the present study, we selected the PTZ-induced seizure model in adult zebrafish to enable a direct comparison of the antiseizure properties of LIN and NER, while simultaneously enhancing the external validity of previously reported findings.

## 2. MATERIALS AND METHODS

### 2.1. Animals

240 adult wild-type zebrafish (6 months old, 300-500 mg) with a similar distribution of males and females were used. Animals were obtained from a local commercial supplier (Distribuidora Flower Pet, Porto Alegre, Brazil) and habituated for at least 2 weeks in 14-L housing tanks (maximum density of 5 animals/L) until the test day. Facility conditions were controlled and settled as ideal for the species: temperature 28 ± 1 °C, pH 7.0 ± 0.5, conductivity 500 uS/cm, luminosity 260 ± 5 lux, and light/dark cycle of 14:10 hours. Fish were fed twice a day with commercial feed (Poytara®, Brazil) and *Artemia salina*. The protocol was approved by the Ethics Committee for Animal Use (CEUA) at the Universidade Federal do Rio Grande do Sul (#36307) and conducted according to the guidelines set by the National Council for the Control of Animal Experimentation (CONCEA).

### 2.2. Chemicals and drugs

Linalool (LIN, CAS 78-70-6, Sigma-Aldrich Product Number L2602), *trans*-nerolidol (NER, CAS 40716-66-3, Sigma-Aldrich Product Number 18143), dimethyl sulfoxide (DMSO, CAS 67-68-5, Sigma-Aldrich Product Number 472301), and pentylenetetrazole (PTZ, CAS 54-95-5, Sigma-Aldrich Product Number P6500) were purchased from Sigma-Aldrich (St. Louis, MO, USA). Diazepam (DZP), used as a positive control, was obtained from União Química Nacional S/A (São Paulo, Brazil). Stock solutions of the test compounds were initially prepared in 10% DMSO and then diluted to ensure a final DMSO concentration of no more than 1%. This concentration has been previously shown not to affect toxicological or behavioral outcomes in zebrafish (17,18). Exposure to 1% DMSO did not induce any significant locomotor changes compared to control groups (see Figures 1–3, Supplementary Material). The concentration range for the tested compounds (4, 40, and 400 μM) was defined based on a logarithmic scale, with the intermediate concentration (40 μM) selected according to previous reports identifying it as an effective anxiolytic dose of LIN in zebrafish (19).

**Figure 1:**
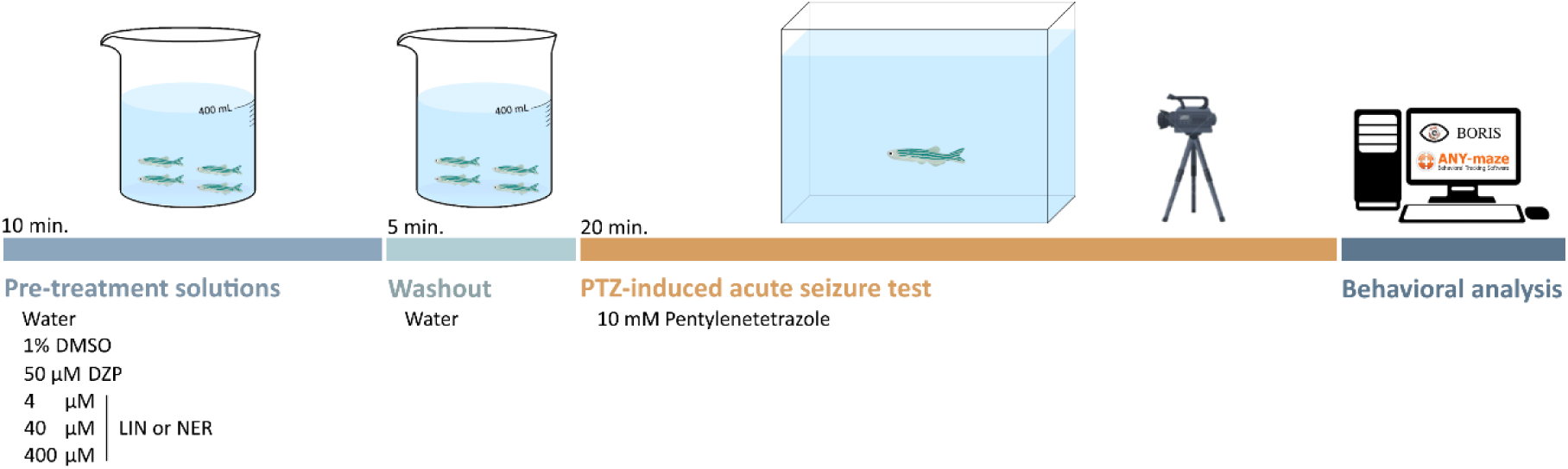
Experimental design showing LIN and NER treatment followed by the acute PTZ-induced seizure test. DMSO = dimethylsulfoxide; DZP = diazepam; LIN = linalool; NER = *trans*-nerolidol.

**Figure 2:**
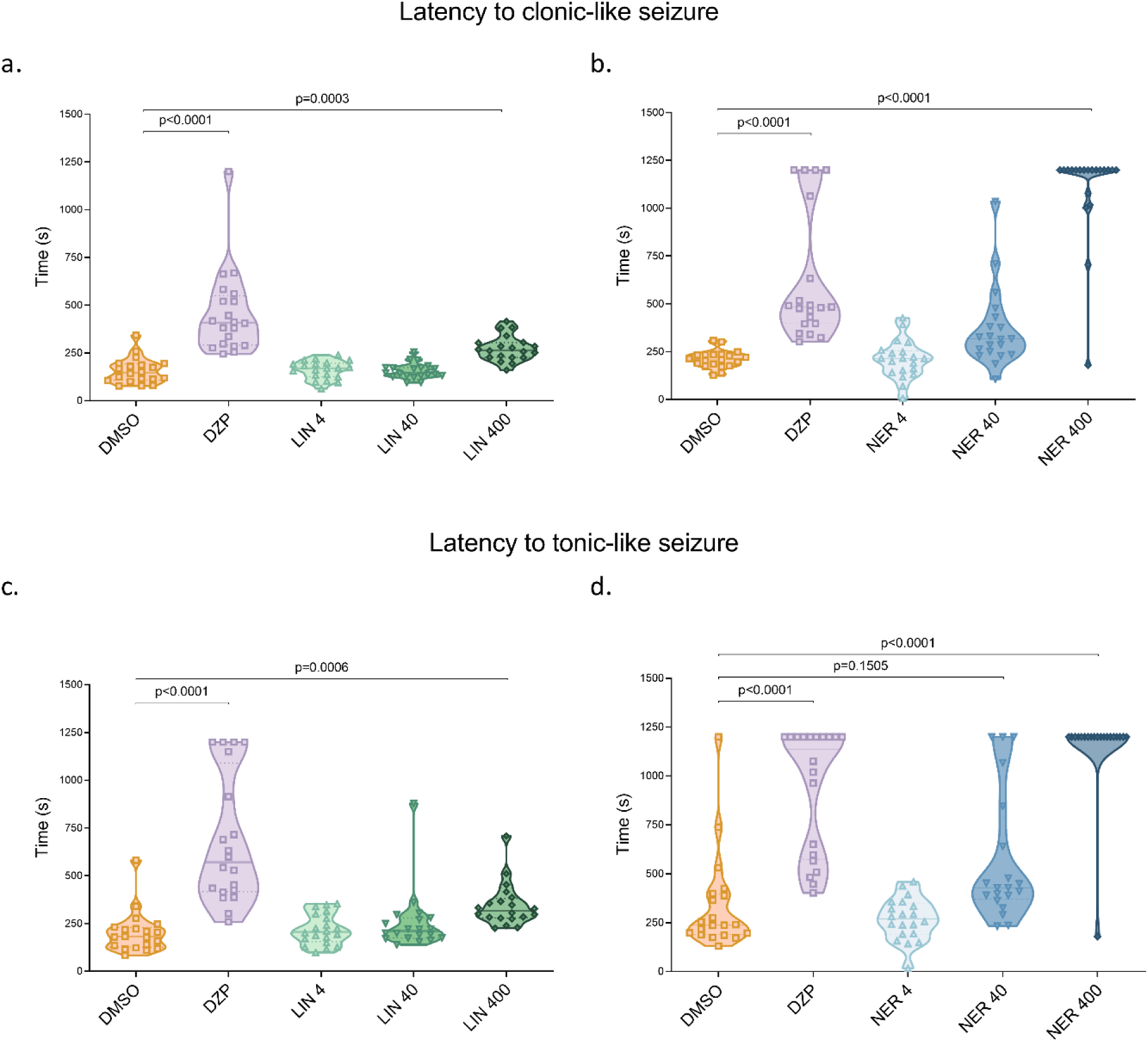
Effects of LIN (4, 40, and 400 μM), NER (4, 40, and 400 μM), and DZP on (a-b) latency to reach clonic-like seizure stage, (c-d) latency to reach tonic-like seizure stage. Data are expressed as median ± interquartile range and were analyzed by Kruskal-Wallis followed by Dunn’s post hoc test. n= 20. DMSO = 1% dimethylsulfoxide; DZP = 50 µM diazepam; LIN = linalool; NER = *trans*-nerolidol.

**Figure 3:**
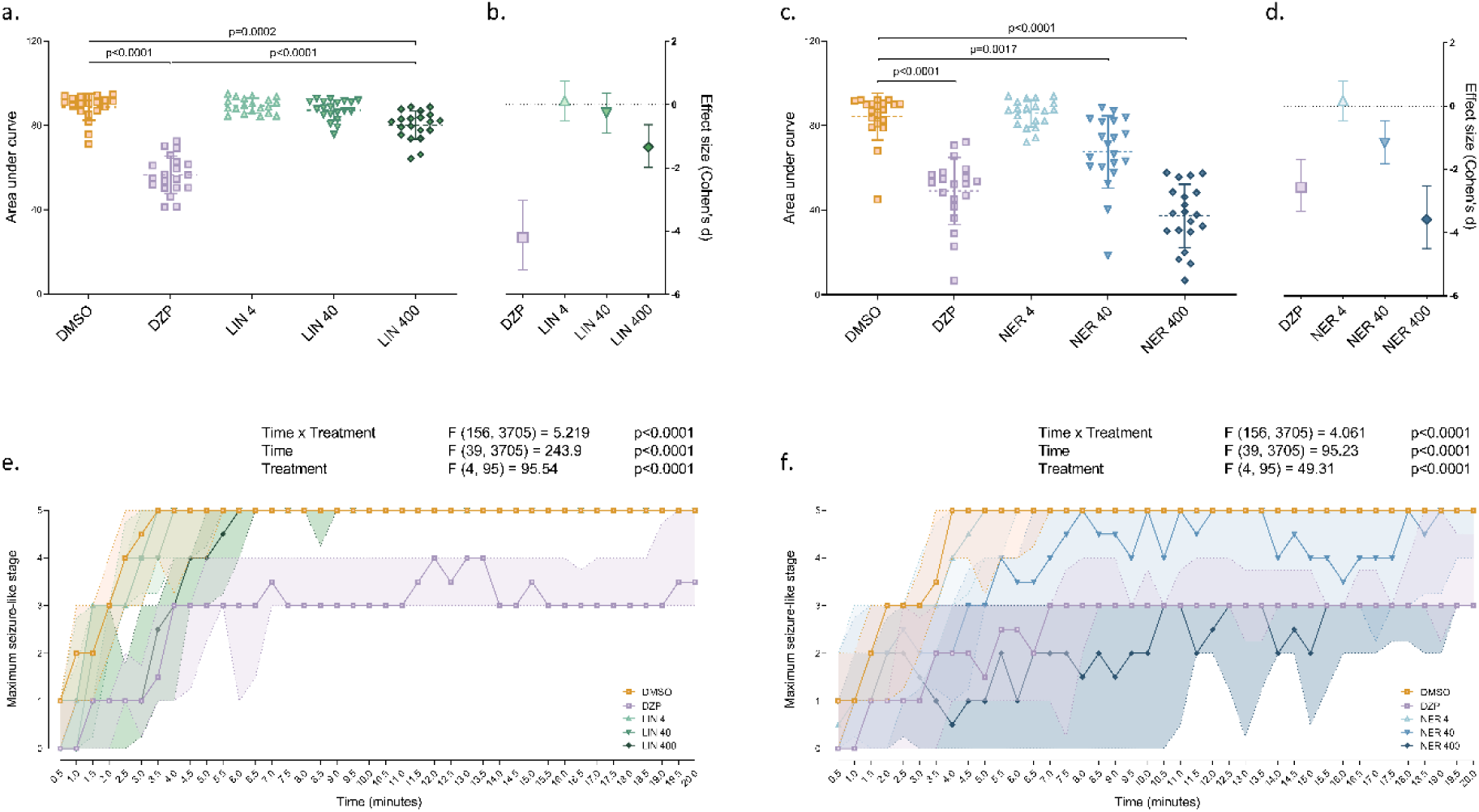
Effects of LIN (4, 40, and 400 μM), NER (4, 40, and 400 μM), and DZP on (a and c) seizure intensity, as calculated by the area under the curve of maximum seizure-like stage across time, (b and d) effect size of seizure intensity, and (e and f) maximum seizure-like stage reached across time. Data are expressed as mean ± S.D. (a and c), standardized mean difference ± 95% confidence interval (b and d), and median ± interquartile range (e and f). Data were analyzed by one-way ANOVA followed by Tukey’s post hoc test (a and c), Cohen’s d effect size test (b and d), and repeated measures two-way ANOVA followed by Tukey’s post hoc test (e and f). n= 20. DMSO = 1% dimethylsulfoxide; DZP = 50 µM diazepam; LIN = linalool; NER = *trans*-nerolidol.

### 2.3. Experimental design

The experimental design was adapted from previous studies (19). Experiments were conducted between 8:00 a.m. and 12:00 p.m. in two batches for each compound. Housing tanks were carefully positioned side by side under similar conditions of lighting, temperature, and elevation.

On the test day, animals were exposed (four fish per beaker) to one of the treatment solutions (400 mL): control (dechlorinated water), vehicle (1% DMSO), positive control (50 µM DZP), or LIN or NER at concentrations of 4, 40, or 400 µM. The treatment lasted 10 minutes, with 20 animals assigned to each experimental group. Allocation to treatment solutions and the test apparatus was randomized using block randomization (random.org), and the source tank was counterbalanced to prevent treatment allocation from a single housing tank. Immediately after treatment exposure, fish were transferred to a washout beaker containing 400 mL of dechlorinated water for 5 minutes. This step was essential to minimize cross-contamination of the PTZ solution, which was not replaced between individual tests. Following the washout period, each fish was individually exposed to the PTZ-induced acute seizure test. This involved placing the animal in a tank (13 cm high × 15 cm long × 10 cm wide) filled with 1 liter of 10 mM PTZ solution, where behavior was recorded for 20 minutes using a Logitech® C920 HD webcam (see Figure 1). Eight tanks containing PTZ were recorded per round. At the end of the test, fish were humanely euthanized by immersion in ice-cold water (0–4 °C) until opercular movement ceased, followed by decapitation, in accordance with the guidelines of the Conselho Nacional de Controle de Experimentação Animal (CONCEA). Although experimenters were not blinded to treatment allocation during the experimental procedures, videos were coded to maintain blinding during behavioral and locomotor analysis.

### 2.4. Seizure-like behavior

Seizure-like behavioral stages were evaluated by researchers blinded to the experimental groups, following established criteria (16): stage 0: Basal locomotor activity; stage 1: Increased swimming activity and opercular movement frequency; stage 2: Whip-like swimming and erratic movements; stage 3: Whirlpool-like behavior and circular swimming; stage 4: Clonic-like seizures with abnormal muscle contractions; stage 5: Tonic-like seizures characterized by loss of posture and sinking to the tank bottom; stage 6: Death. Each fish was scored individually, and researchers recorded the latency to reach clonic (Stage 4) and tonic-like (Stage 5) seizures, as well as the maximum seizure-like stage reached every 30 seconds over a 20-minute period. To maintain blinding throughout the analysis, experimental groups were manually coded by a third researcher until the data assessment was complete. Seizure intensity was calculated as the area under the curve (AUC) of the maximum seizure stage reached over time. Behavioral analyses were conducted using BORIS® version 7.7.3 (Behavioral Observation Research Interactive Software) (20).

### 2.5. Locomotion

Locomotor behavior during the acute seizure tests was analyzed using ANY-maze™ v. 7.4 tracking software (Stoelting Co., Wood Dale, IL, USA) by researchers blinded to the experimental groups. Group identities were encoded within the software until the completion of the analysis to ensure unbiased outcome assessments. The parameters evaluated included total distance traveled, mean swimming speed during the test, and distance traveled over time (measured in meters per 30-second bins).

### 2.6. Statistical analysis

The sample size was calculated using G*Power 3.1.9.6 software, with seizure intensity defined as the primary outcome. A fixed-effects ANOVA model was applied using the following parameters: α = 0.05, statistical power = 0.95, and effect size = 0.4, considering six experimental groups. This yielded a total required sample size of 114 animals (n = 19 per group). To account for potential data loss due to tracking errors, one additional animal was included per group (n = 20). The full dataset from this study is available in Open Science Framework (OSF) repository at https://osf.io/8qg4r/ (21). All animals were included in the final analysis. The normality of variance was assessed using the D’Agostino & Pearson test. Latencies to reach clonic and tonic-like seizure stages were analyzed using Kruskal–Wallis tests followed by Dunn’s post hoc analysis. Seizure-like stages over time and distance traveled over time were analyzed using repeated measures two-way ANOVA (Factor 1: time; Factor 2: drug), followed by Tukey’s post hoc test. Seizure intensity, total distance traveled, and mean swimming speed were analyzed using one-way ANOVA, followed by Tukey’s post hoc test. Effect sizes for seizure intensity were calculated using Cohen’s d for independent samples via an online calculator (https://www.cem.org/effect-size-calculator). Parametric data are presented as mean ± standard deviation (SD), non-parametric data as median ± interquartile range (IQR), and effect sizes as standardized mean difference ± 95% confidence interval (CI). Significance was set at p < 0.05. All statistical analyses and graph generation were performed using GraphPad Prism version 8.0.1.

## 3. RESULTS

No significant differences were observed between the 1% DMSO and dechlorinated water control groups in terms of latency to the highest seizure-like stages or seizure intensity (Supplementary Figures 1-3). Therefore, all data are expressed relative to the 1% DMSO group, which reflects the highest DMSO concentration used in the treatment solutions (400 µM LIN and NER).

As expected, DZP significantly increased the latency to reach both clonic- and tonic-like seizure stages in the LIN and NER experiments (Figures 2a-d; Kruskal-Wallis test followed by Dunn’s post hoc test, H(4)=64.17 and H(4)=55.01, p<0.0001, and H(4)=67.61 and H(4)=67.61, p<0.0001, respectively). LIN at the highest concentration (400 µM) significantly increased the latency to reach clonic- and tonic-like seizure stages (Figures 2a and 2c; H(4)=64.17, p=0.0003, and H(4)=55.01, p=0.0006). NER at the highest concentration (400 µM) significantly increased the latency to the clonic-like seizure stage (Figure 2b; H(4)=67.61, p<0.0001), while both 40 µM and 400 µM increased the latency to the tonic-like stage (Figure 2d; H(4)=61.91, p=0.1505 and p<0.0001, respectively).

DZP exposure reduced seizure intensity, quantified as the area under the curve (AUC), in both experiments (Figures 3a and 3c; One-way ANOVA followed by Tukey’s post hoc test, F (4, 95) = 96.72, p<0.0001, and F (4, 95) = 49.38, p<0.0001). The maximum seizure-like stage reached at each 30-second interval is shown in Figures 3e (LIN) and 3f (NER). Effect size analysis revealed a large biological effect of DZP on seizure intensity attenuation in both experiments (Cohen’s d = 4.20 and 2.58, respectively; Figures 3b and 3d). Linalool at 400 µM also significantly reduced seizure intensity compared to control (Figure 3a; F (4, 95) = 96.72, p=0.0002). However, LIN’s effect was significantly smaller than that of DZP (Figure 3a; F (4, 95) = 96.72, p<0.0001), although it still showed a large biological effect size (Cohen’s d = 1.34; Figure 3b). The same NER concentrations (40 and 400 µM) that increased latency also significantly reduced seizure intensity (Figure 3c; F (4, 95) = 49.38, p=0.0017 and p<0.0001, respectively), with large effect sizes (Cohen’s d = 1.17 and 3.58; Figure 3d). Notably, the effect size for 400 µM NER was larger than that for DZP (Cohen’s d = 3.58 vs. 2.58; Figure 3d). Maximum seizure-like stage increased across time in LIN and NER experiments (Figures 3e and 3f; repeated-measures two-way ANOVA followed by Tukey’s post hoc test; Time Factor (LIN experiment): F (39, 3705) = 243.9, p<0.0001; Time Factor (NER experiment): F (39, 3705) = 95.23, p<0.0001). Drug treatments (DZP, LIN, and NER) decreased the maximum seizure-like stage reached during the PTZ exposure (Figure 3e and 3f; repeated-measures two-way ANOVA, Drug Factor (LIN experiment): F(4, 95) = 95.54; p<0.0001; Drug Factor (NER experiment): F(4, 95) = 49.31; p<0.0001). Finally, an interaction between these factors was observed in both LIN and NER experiments (Figure 3e and 3f; Interaction Factor (LIN experiment): F (156,3705) = 5.219; p<0.0001; Interaction Factor (NER experiment): F (156,3705) = 4.061; p<0.0001). Post hoc comparisons showed that 40 µM LIN differed from DMSO at the first minute-bin, 400 µM LIN from 1–5.5 minutes, and DZP across the entire LIN experiment. In the NER experiment, 40 µM NER differed from DMSO at 3.5–7 and 16–17 minutes, 400 µM NER from 3.5 minutes to the end, and DZP from 1.5 minutes to the end.

Neither DZP nor LIN altered total distance traveled or mean swimming speed (DZP: Figures 4a-4d, LIN: Figure 4a and 4c; one-way ANOVA, F (4, 95) = 1.346). However, NER at 40 µM increased both total distance traveled and mean swimming speed (Figures 4b and 4d; one-way ANOVA followed by Tukey’s post hoc test, F (4, 95) = 2.598, p=0.0262, and p=0.0239, respectively). Distance traveled decreased over time in a time-dependent manner in both experiments (Figures 5a and 5b; repeated-measures two-way ANOVA followed by Tukey’s post hoc test; Time Factor (LIN experiment): F (39, 3705) = 10.69, p<0.0001; Time Factor (NER experiment): F (39, 3705) = 1.488, p=0.0263). The drug treatment factor significantly influenced distance traveled in the 40 µM NER group (Figure 5b; repeated-measures two-way ANOVA, Drug Factor: F(4, 95) = 2.598, p=0.0410). A significant interaction between time and drug exposure was observed for both LIN (Figure 5a; repeated-measures Two-way ANOVA, Interaction Factor: F (156, 3705) = 1.284, p=0.0112) and NER (Figure 5b; repeated-measures Two-way ANOVA, Interaction Factor: F (156, 3705) = 1.371, p=0.0019). Post hoc comparisons showed that 4 and 40 µM LIN differed from DMSO only at the 6.5-minute bin, and 400 µM LIN at 4–5 minutes. In the NER experiment, 40 µM NER differed from DMSO only at the 5-minute bin, 400 µM NER at 0.5–1 minute, and DZP at 7.5–8 minutes.

**Figure 4:**
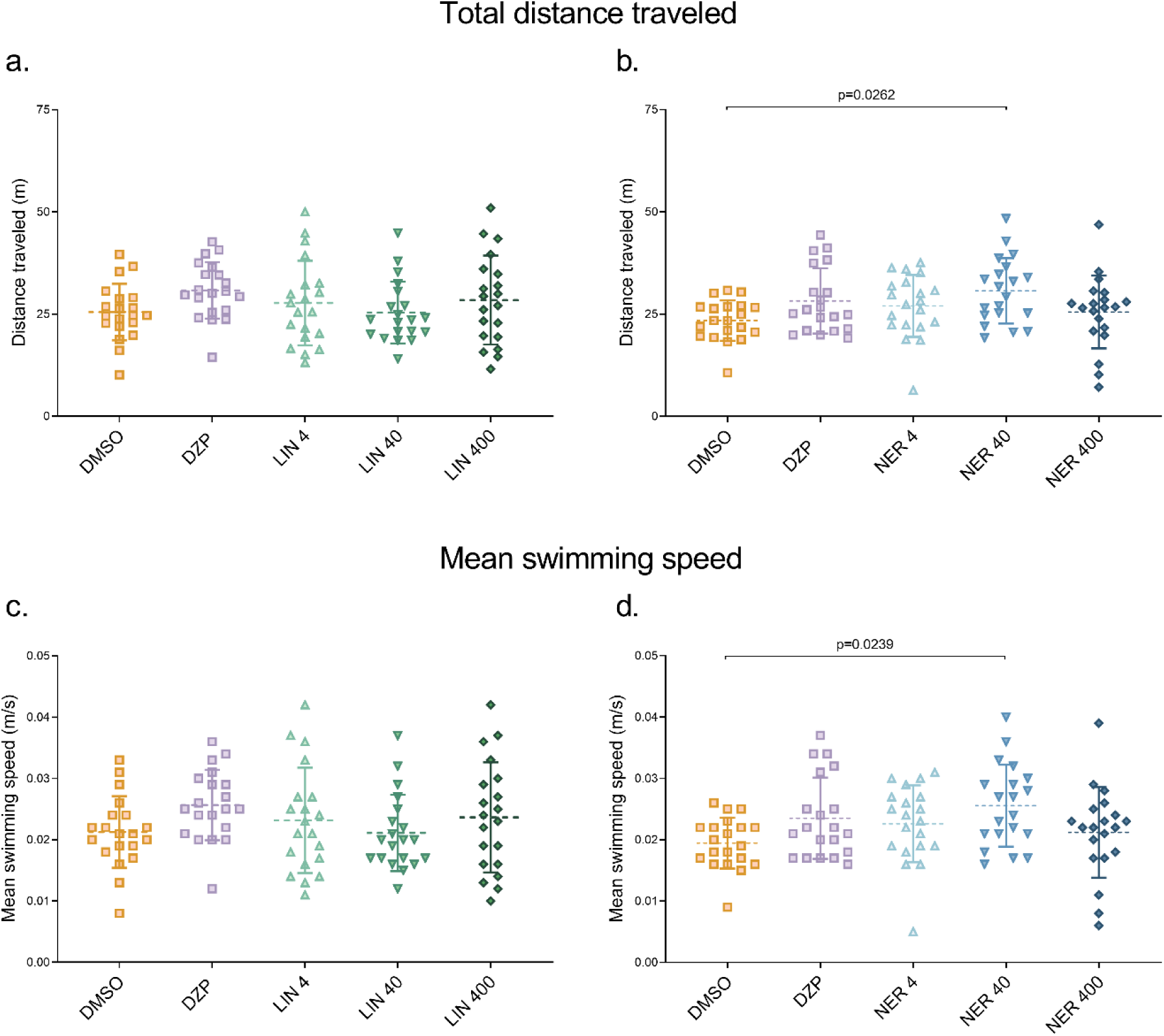
Effects of LIN (4, 40, and 400 μM), NER (4, 40, and 400 μM), and DZP on (a-b) total distance traveled, and (c-d) mean swimming speed. Data are expressed as mean ± S.D. and were analyzed by one-way ANOVA followed by Tukey’s post hoc. n= 20. DMSO = 1% dimethylsulfoxide; DZP = 50 µM diazepam; LIN = linalool; NER = *trans*-nerolidol.

**Figure 5:**
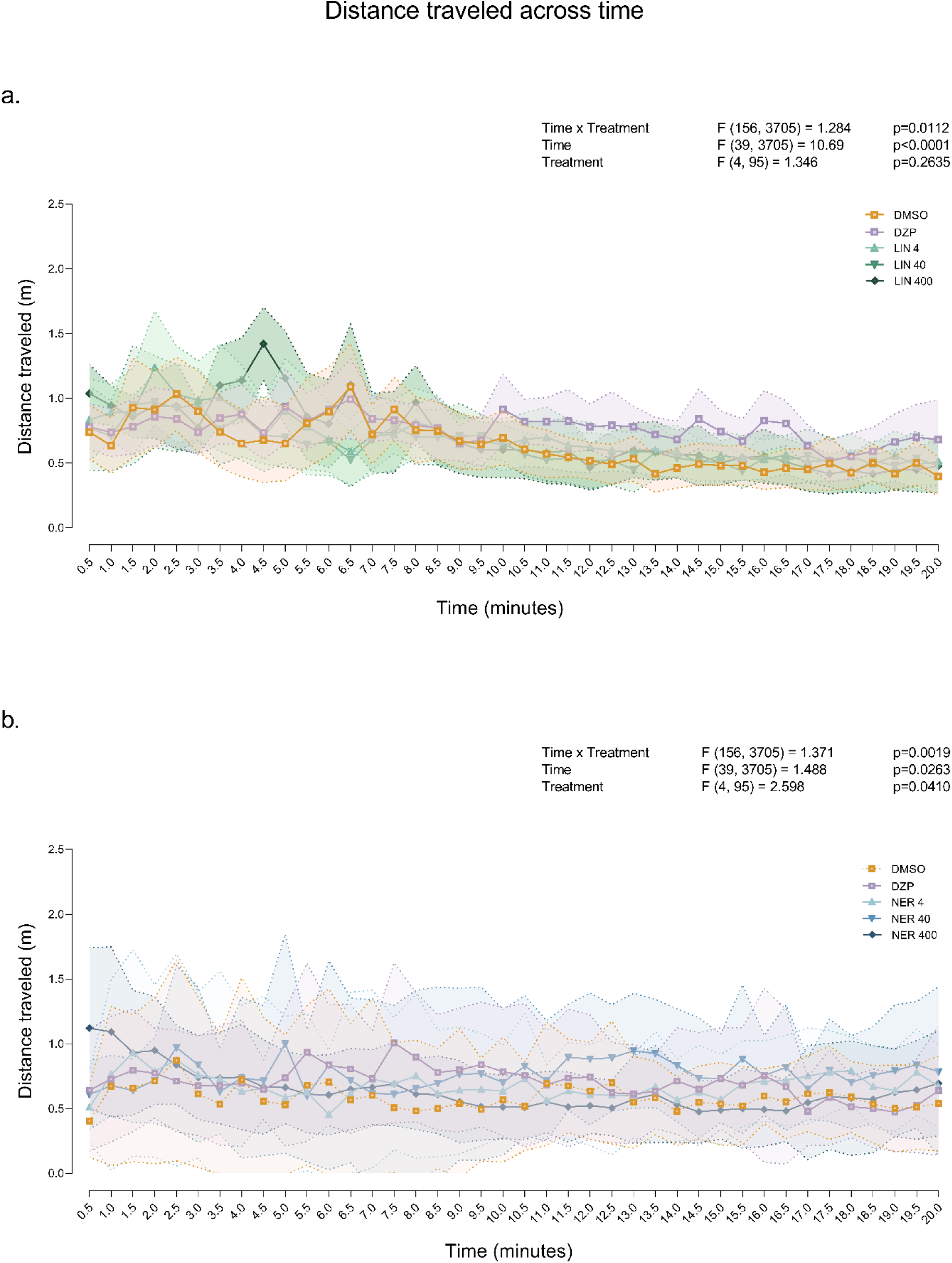
Effects of LIN (4, 40, and 400 μM), NER (4, 40, and 400 μM), and DZP on (a-b) distance traveled across time. Data are expressed as mean ± S.D. and were analyzed by repeated measures two-way ANOVA. n= 20. DMSO = 1% dimethylsulfoxide; DZP = 50 µM diazepam; LIN = linalool; NER = *trans*-nerolidol.

## 4. DISCUSSION

In this study, we demonstrated the antiseizure properties of linalool (LIN) and *trans*-nerolidol (NER) using a PTZ-induced acute seizure model in adult zebrafish. Acute exposure to both compounds produced large effect sizes, significantly increasing the latency to clonic- and tonic-like seizure stages and reducing seizure intensity.

Linalool (LIN) is a monoterpene synthesized via the mevalonate pathway, which plays a key role in plant metabolism by converting acetyl-CoA into isoprenoids such as cholesterol and steroid hormones (10,22,23). LIN has demonstrated antiseizure properties in both rats and mice, increasing seizure latency and reducing the duration of seizure-like behaviors in models involving glutamatergic and GABAergic pathways (24–27). However, in zebrafish larvae, LIN was reported to be inactive against both chemically induced and SCN1a-mutant seizure models at concentrations ranging from 0.3 to 4 µM (28). The antiseizure effects observed in our study are consistent with the rodent data, with efficacy seen at a LIN concentration of 400 µM—approximately 100 times higher than those tested by Thornton and colleagues, where 4 µM LIN was also ineffective.

*Trans*-nerolidol (NER) is the *trans* isomer of a naturally occurring sesquiterpene, biosynthesized via the 2-C-methyl-D-erythritol 4-phosphate (MEP) pathway. This mevalonate-independent route also plays a crucial role in isoprenoid biosynthesis by converting pyruvate, serving as an alternative to the mevalonate pathway (29–31). Kaur and colleagues (32) investigated the effects of chronic NER administration in a PTZ-induced kindling model in mice and reported a reduction in seizure severity. Similarly, Neroli essential oil, which contains high concentrations of NER, has been shown to prevent clonic- and tonic-like seizures in PTZ and maximal electroshock-induced seizure models in mice (9). In the present study, we demonstrate for the first time the antiseizure effects of NER in a zebrafish seizure model. Acute exposure to NER increased the latency to the most severe seizure-like stages and significantly reduced seizure intensity. Notably, the effect size observed at 400 µM NER was larger than that of 50 µM diazepam. This may be attributed to NER’s higher lipophilicity, stemming from its longer carbon chain and lower molecular weight (1), which may enhance its bioavailability and ability to cross the blood-brain barrier more readily than diazepam.

The underlying mechanisms of the antiseizure effects observed in this study have yet to be clarified. LIN has been shown to potentiate GABAergic currents through the GABA_A_ receptor α1β2γ2 subunits in vivo (33–35), and to bind to NMDA receptors and serotonin transporters in a dose-dependent manner (36). Flumazenil prevented the antiseizure effects of an essential oil (EO) containing a high concentration of NER in mice (9). Additionally, in an antinociceptive study, the effects of NER were reversed by the GABA_A_ antagonist bicuculline (37). Therefore, it is likely that the antiseizure activity of both compounds involves interaction with the benzodiazepine binding site of GABA_A_ receptors. Moreover, since the pathophysiology of epilepsy also includes neuroinflammation (38) and oxidative stress (39), and given that both LIN and NER have demonstrated antioxidant and anti-inflammatory properties (40–43), the beneficial effects of these compounds in epilepsy syndromes might not be limited to the attenuation of PTZ-induced seizures (44,45). PTZ-induced changes in locomotion include increased distance traveled and swimming speed in both larval and adult zebrafish (3,46). As most antiseizure drugs reduce locomotion in zebrafish (46), surprisingly, 40 µM *trans*-nerolidol increased both distance traveled and swimming speed — unlike at similar doses that minimize PTZ-induced seizures. The biological significance of this finding remains unclear and warrants further investigation.

Although PTZ is the most commonly used chemical seizure-inducing agent model in zebrafish drug screening (14), it is an acute seizure model that does not replicate the comorbidities and neurocognitive impairments observed in clinical epilepsy. Therefore, despite the positive results presented here, this constitutes a limitation of the study. The potential therapeutic value of these compounds cannot be fully assessed without additional data from chronic epilepsy models, such as kindling or genetic models, which more closely resemble patients with spontaneous and recurrent seizures (38,47).

## 5. CONCLUSION

Linalool is a monoterpene alcohol with a smaller, more polar structure, while *trans*-nerolidol is a sesquiterpene alcohol with a larger, more lipophilic carbon backbone. The increased lipophilicity of *trans*-nerolidol may contribute to its prolonged tissue distribution and distinct pharmacological profile. These structural differences likely influence their receptor interactions and bioavailability. This is the first study to report the behavioral effects of these compounds in adult zebrafish within an antiseizure context. Further research is needed to better characterize linalool and *trans*-nerolidol effects, particularly through electrophysiological and pharmacodynamic assessments.

## List of abbreviations

AUC: Area under the curve
DMSO: Dimethylsulfoxide
DZP: Diazepam
EOs: Essential oils
LIN: Linalool
NER: *trans*-nerolidol
PTZ: Pentylenetetrazole

## AUTHORS CONTRIBUTION

Conceptualization: D.S.N., E.E., A.P.H., and A.P. Data curation: A.L.S. and L.M.B. Formal analysis: A.L.S., L.M.B., M.G.-L. and M.E.C. Funding acquisition: A.P. Investigation: A.L.S., L.M.B., R.C., C.G.R., and M.G.-L. Methodology: A.L.S., L.M.B., R.C., C.G.R., and M.G.-L. Project administration: A.P. Supervision: A.P. Writing - original draft: A.L.S., L.M.B., and A.P. Writing - review & editing: L.M.B., A.L.S., M.G.-L., R.C., C.G.R., D.S.N., E.E., A.P.H., M.E.C., and A.P.

## DATA AVAILABILITY

The full dataset from this study is available in Open Science Framework (OSF) repository at https://osf.io/8qg4r/.

## SUPPLEMENTARY MATERIAL

**Supplementary Figure 1:**
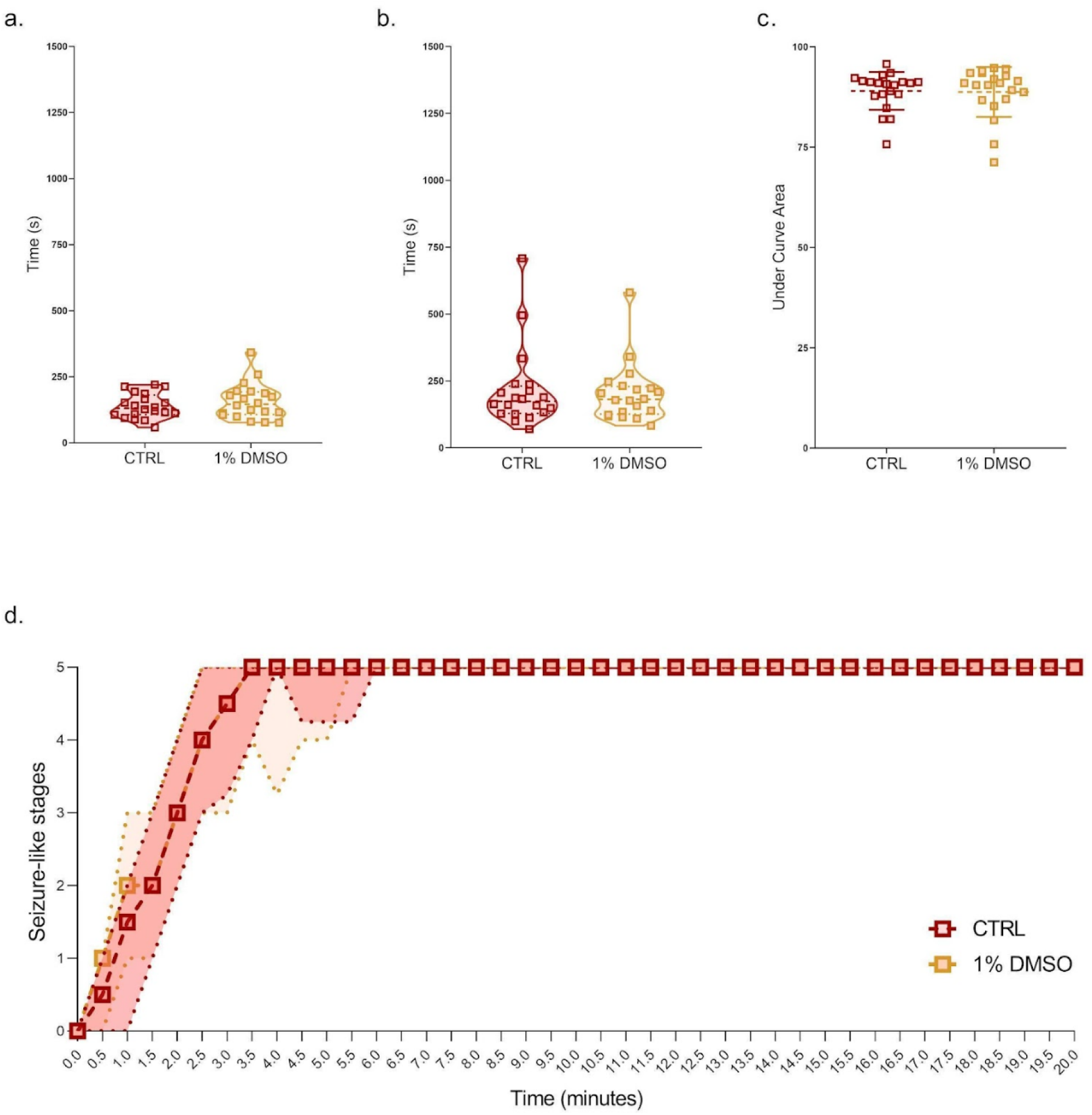
Linalool experiment: effects of 1% DMSO (vehicle) on (a) latency to clonic-like seizure stage (score onset 4), (b) latency to tonic-like seizure stage (score onset 5), (c) seizure intensity and (d) seizure-like stages scored across time. Data were analyzed for normality by D’agostino & Pearson test, and for variance differences between groups by T-test (Fig. 1a and 1b) and Mann-Whitney test (Fig. 1c) according to data distribution. n=20.

**Supplementary Figure 2:**
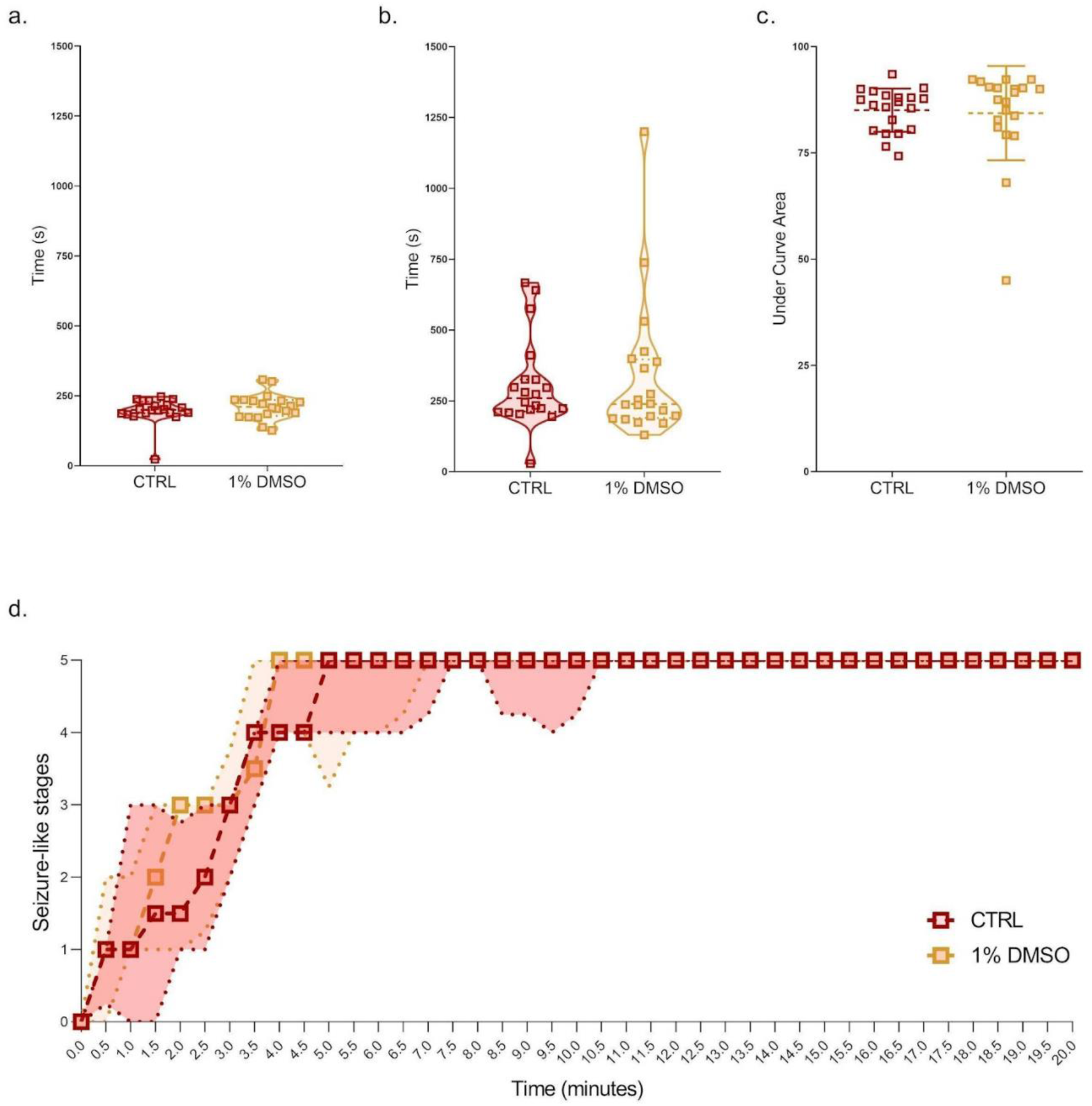
*Trans*-nerolidol experiment: effects of 1% DMSO (vehicle) on (a) latency to clonic-like seizure stage (score onset 4), (b) latency to tonic-like seizure stage (score onset 5), (c) seizure intensity and (d) seizure-like stages scored across time. Data were analyzed for normality by D’agostino & Pearson test, and for variance differences between groups by T-test (Fig. 2a and 2b) and Mann-Whitney test (Fig. 2c) according to data distribution. n=20.

**Supplementary Figure 3:**
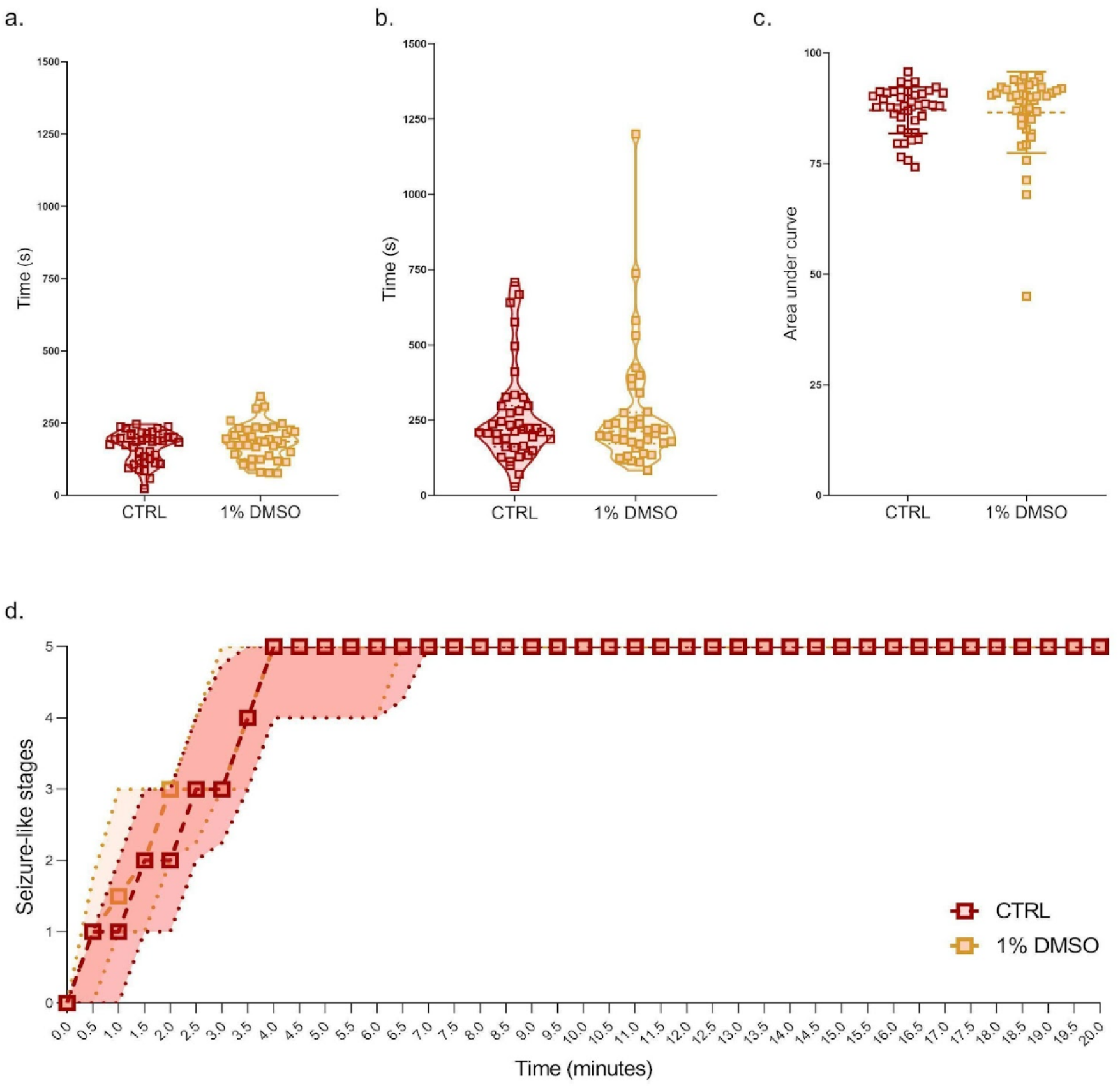
Linalool and *trans*-nerolidol experiments cumulative data - Effects of 1% DMSO (vehicle) on (a) latency to clonic-like seizure stage (score onset 4), (b) latency to tonic-like seizure stage (score onset 5), (c) seizure intensity and (d) seizure-like stages scored across time. Data were analyzed for normality by D’agostino & Pearson test, and for variance differences between groups by T-test (Fig. 3a and 3b) and Mann-Whitney test (Fig. 3c) according to data distribution. n=40.

## REFERENCES

1. Elaine Elisabetsky, Domingos S. Nunes. Central Nervous System Effects of Essential Oil Compounds. In: Handbook of Essential Oils. 3rd ed. London: CRC Press; 2020. p. 303–44.

2. Wang ZJ, Heinbockel T. Essential Oils and Their Constituents Targeting the GABAergic System and Sodium Channels as Treatment of Neurological Diseases. Molecules. 2018 May;23(5):1061.

3. Lee Y, Kim D, Kim YH, Lee H, Lee CJ. Improvement of pentylenetetrazol-induced learning deficits by valproic acid in the adult zebrafish. Eur J Pharmacol. 2010;643(2):225–31.

4. Nóbrega de Almeida R, Agra M de F, Negromonte Souto Maior F, De Sousa DP. Essential Oils and Their Constituents: Anticonvulsant Activity. Molecules. 2011 Mar;16(3):2726–42.

5. Paul A, Cox PA. An Ethnobotanical Survey of the Uses for Citrus aurantium (Rutaceae) in Haiti. Econ Bot. 1995;49(3):249–56.

6. Carvalho-Freitas MIR, Costa M. Anxiolytic and Sedative Effects of Extracts and Essential Oil from *Citrus aurantium* L. Biol Pharm Bull. 2002;25(12):1629–33.

7. Elisabetsky E, Brum L. Linalool as active component of traditional remedies: anticonvulsant properties and mechanisms of action. Curare. 2003 Jan 1;26:45–52.

8. Boleti AP de A, Frihling BEF, e Silva PS, Cardoso PH de O, de Moraes LFRN, Rodrigues TAA, et al. Biochemical aspects and therapeutic mechanisms of cannabidiol in epilepsy. Neurosci Biobehav Rev. 2022;132:1214–28.

9. Azanchi T, Shafaroodi H, Asgarpanah J. Anticonvulsant activity of Citrus aurantium blossom essential oil (neroli): involvment of the GABAergic system. Nat Prod Commun. 2014 Nov;9(11):1615–8.

10. Cláudia Maria Oliveira Simões, Eloir Paulo Schenkel, João Carlos Palazzo de Mello, Lilian Auler Mentz, Pedro Ros Petrovick. Farmacognosia: Do Produto Natural ao Medicamento. 1st ed. Artmed; 2016.

11. Klopell FC, Lemos M, Sousa JPB, Comunello E, Maistro EL, Bastos JK, et al. Nerolidol, an Antiulcer Constituent from the Essential Oil of Baccharis dracunculifolia DC (Asteraceae). Z Für Naturforschung C. 2007 Aug 1;62(7–8):537–42.

12. Pokajewicz K, Białoń M, Svydenko L, Fedin R, Hudz N. Chemical Composition of the Essential Oil of the New Cultivars of Lavandula angustifolia Mill. Bred in Ukraine. Molecules. 2021 Sep 18;26(18):5681.

13. Queiroga C, Fukai A, Marsaioli A. Composition of the Essential Oil of Vassoura. J Braz Chem Soc. 1990 Jan 1;1:105–9.

14. Chitolina R, Gallas-Lopes M, Reis CG, Benvenutti R, Stahlhofer-Buss T, Calcagnotto ME, et al. Chemically-induced epileptic seizures in zebrafish: A systematic review. Epilepsy Res. 2023 Nov 1;197:107236.

15. Baraban SC, Taylor MR, Castro PA, Baier H. Pentylenetetrazole induced changes in zebrafish behavior, neural activity and c-fos expression. Neuroscience. 2005;131(3):759–68.

16. Mussulini BHM, Leite CE, Zenki KC, Moro L, Baggio S, Rico EP, et al. Seizures Induced by Pentylenetetrazole in the Adult Zebrafish: A Detailed Behavioral Characterization. PLOS ONE. 2013 Jan 21;8(1):e54515.

17. Sackerman J, Donegan JJ, Cunningham CS, Nguyen NN, Lawless K, Long A, et al. Zebrafish Behavior in Novel Environments: Effects of Acute Exposure to Anxiolytic Compounds and Choice of Danio rerio Line. Int J Comp Psychol ISCP Spons Int Soc Comp Psychol Univ Calabr. 2010 Jan 1;23(1):43–61.

18. Hoyberghs J, Bars C, Ayuso M, Van Ginneken C, Foubert K, Van Cruchten S. DMSO Concentrations up to 1% are Safe to be Used in the Zebrafish Embryo Developmental Toxicity Assay. Front Toxicol. 2021;3:804033.

19. Szaszkiewicz J, Leigh S, Hamilton TJ. Robust behavioural effects in response to acute, but not repeated, terpene administration in Zebrafish (Danio rerio). Sci Rep. 2021 Sep 28;11(1):19214.

20. Friard O, Gamba M. BORIS: a free, versatile open-source event-logging software for video/audio coding and live observations. Methods Ecol Evol. 2016;7(11):1325–30.

21. Silva AL, Bastos LM, Chitolina R, Reis CG, Gallas-Lopes M, Buss TS, et al. Linalool and trans-nerolidol effects in pentylenetetrazole-induced seizure model in zebrafish. 2023 Aug 23 [cited 2025 Jan 13]; Available from: https://osf.io/8qg4r/

22. Buhaescu I, Izzedine H. Mevalonate pathway: a review of clinical and therapeutical implications. Clin Biochem. 2007 Jun;40(9–10):575–84.

23. Liao P, Hemmerlin A, Bach TJ, Chye ML. The potential of the mevalonate pathway for enhanced isoprenoid production. Biotechnol Adv. 2016;34(5):697–713.

24. Elisabetsky E, Brum LF, Souza DO. Anticonvulsant properties of linalool in glutamate-related seizure models. Phytomedicine Int J Phytother Phytopharm. 1999 May;6(2):107–13.

25. Coelho de Souza GP, Elisabetsky E, Nunes DS, Rabelo SKL, Nascimento da Silva M. Anticonvulsant properties of γ-decanolactone in mice. J Ethnopharmacol. 1997;58(3):175–81.

26. Almeida ER de, Rafael KR de O, Couto GBL, Ishigami ABM. Anxiolytic and Anticonvulsant Effects on Mice of Flavonoids, Linalool, and alpha-Tocopherol Presents in the Extract of Leaves of Cissus sicyoides L. (Vitaceae). BioMed Res Int. 2009 Mar 12;2009:e274740.

27. Bahr TA, Rodriguez D, Beaumont C, Allred K. The Effects of Various Essential Oils on Epilepsy and Acute Seizure: A Systematic Review. Evid Based Complement Alternat Med. 2019 May 22;2019:e6216745.

28. Thornton C, Dickson KE, Carty DR, Ashpole NM, Willett KL. Cannabis constituents reduce seizure behavior in chemically-induced and scn1a-mutant zebrafish. Epilepsy Behav EB. 2020 Sep;110:107152.

29. Eisenreich W, Bacher A, Arigoni D, Rohdich F. Biosynthesis of isoprenoids via the non-mevalonate pathway. Cell Mol Life Sci CMLS. 2004 Jun;61(12):1401–26.

30. Hunter WN. The Non-mevalonate Pathway of Isoprenoid Precursor Biosynthesis*. J Biol Chem. 2007 Jul 27;282(30):21573–7.

31. Rohmer M. The discovery of a mevalonate-independent pathway for isoprenoid biosynthesis in bacteria, algae and higher plants. Nat Prod Rep. 1999 Oct;16(5):565–74.

32. Kaur D, Pahwa P, Goel RK. Protective Effect of Nerolidol Against Pentylenetetrazol-Induced Kindling, Oxidative Stress and Associated Behavioral Comorbidities in Mice. Neurochem Res. 2016 Nov 1;41(11):2859–67.

33. Hossain SJ, Hamamoto K, Aoshima H, Hara Y. Effects of Tea Components on the Response of GABAA Receptors Expressed in Xenopus Oocytes. J Agric Food Chem. 2002 Jul 1;50(14):3954–60.

34. Kessler A, Sahin-Nadeem H, Lummis SCR, Weigel I, Pischetsrieder M, Buettner A, et al. GABAA receptor modulation by terpenoids from Sideritis extracts. Mol Nutr Food Res. 2014;58(4):851– 62.

35. Milanos S, Elsharif SA, Janzen D, Buettner A, Villmann C. Metabolic Products of Linalool and Modulation of GABAA Receptors. Front Chem [Internet]. 2017 [cited 2024 Jan 30];5. Available from: https://www.frontiersin.org/articles/10.3389/fchem.2017.00046

36. López V, Nielsen B, Solas M, Ramírez MJ, Jäger AK. Exploring Pharmacological Mechanisms of Lavender (Lavandula angustifolia) Essential Oil on Central Nervous System Targets. Front Pharmacol [Internet]. 2017 [cited 2024 Jan 30];8. Available from: https://www.frontiersin.org/articles/10.3389/fphar.2017.00280

37. Fonsêca DV, Salgado PRR, de Carvalho FL, Salvadori MGSS, Penha ARS, Leite FC, et al. Nerolidol exhibits antinociceptive and anti-inflammatory activity: involvement of the GABAergic system and proinflammatory cytokines. Fundam Clin Pharmacol. 2016;30(1):14–22.

38. Devinsky O, Vezzani A, O’Brien TJ, Jette N, Scheffer IE, de Curtis M, et al. Epilepsy. Nat Rev Dis Primer. 2018 May 3;4(1):1–24.

39. Ramos-Riera KP, Pérez-Severiano F, López-Meraz ML. Oxidative stress: a common imbalance in diabetes and epilepsy. Metab Brain Dis. 2023 Mar 1;38(3):767–82.

40. Balakrishnan V, Ganapathy S, Veerasamy V, Duraisamy R, Sathiavakoo VA, Krishnamoorthy V, et al. Anticancer and antioxidant profiling effects of Nerolidol against DMBA induced oral experimental carcinogenesis. J Biochem Mol Toxicol. 2022;36(6):e23029.

41. de Moura DF, Rocha TA, de Melo Barros D, da Silva MM, dos Santos Santana M, Neta BM, et al. Evaluation of the antioxidant, antibacterial, and antibiofilm activity of the sesquiterpene nerolidol. Arch Microbiol. 2021 Sep 1;203(7):4303–11.

42. Lima DKS, Ballico LJ, Rocha Lapa F, Gonçalves HP, de Souza LM, Iacomini M, et al. Evaluation of the antinociceptive, anti-inflammatory and gastric antiulcer activities of the essential oil from Piper aleyreanum C.DC in rodents. J Ethnopharmacol. 2012;142(1):274–82.

43. Pinheiro BG, Silva ASB, Souza GEP, Figueiredo JG, Cunha FQ, Lahlou S, et al. Chemical composition, antinociceptive and anti-inflammatory effects in rodents of the essential oil of Peperomia serpens (Sw.) Loud. J Ethnopharmacol. 2011;138(2):479–86.

44. Hosseini M, Boskabady MH, Khazdair MR. Neuroprotective effects of Coriandrum sativum and its constituent, linalool: A review. Avicenna J Phytomedicine. 2021;11(5):436–50.

45. Linck VM, da Silva AL, Figueiró M, Caramão EB, Moreno PRH, Elisabetsky E. Effects of inhaled Linalool in anxiety, social interaction and aggressive behavior in mice. Phytomedicine. 2010 Jul 1;17(8):679–83.

46. Afrikanova T, Serruys ASK, Buenafe OEM, Clinckers R, Smolders I, Witte PAM de, et al. Validation of the Zebrafish Pentylenetetrazol Seizure Model: Locomotor versus Electrographic Responses to Antiepileptic Drugs. PLOS ONE. 2013 Jan 14;8(1):e54166.

47. Löscher W. Animal Models of Seizures and Epilepsy: Past, Present, and Future Role for the Discovery of Antiseizure Drugs. Neurochem Res. 2017 Jul 1;42(7):1873–88.

